# ARF6 is a host factor for SARS-CoV-2 infection *in vitro*

**DOI:** 10.1101/2022.06.09.495482

**Authors:** C. Mirabelli, E.J. Sherman, J. Bragazzi Cunha, J.W. Wotring, J. El Saghir, J. Harder, M. Kretzler, J.Z. Sexton, B.T. Emmer, C.E. Wobus

## Abstract

SARS-CoV-2 is a newly emerged beta-coronavirus that enter cells via two routes, direct fusion at the plasma membrane or endocytosis followed by fusion with the late endosome/lysosome. While the viral receptor, ACE2, multiple entry factors, and the mechanism of fusion of the virus at the plasma membrane have been extensively investigated, viral entry via the endocytic pathway is less understood. By using a human hepatocarcinoma cell line, Huh-7, which is resistant to the antiviral action of the TMPRSS2 inhibitor camostat, we discovered that SARS-CoV-2 entry is not dependent on dynamin but dependent on cholesterol. ADP-ribosylation factor 6 (ARF6) has been described as a host factor for SARS-CoV-2 replication and it is involved in the entry and infection of several pathogenic viruses. Using CRISPR-Cas9 genetic deletion, we observed that ARF6 is important for SARS-CoV-2 uptake and infection in Huh-7. This finding was corroborated using a pharmacologic inhibitor, whereby the ARF6 inhibitor NAV-2729 showed a dose-dependent inhibition of viral infection. Importantly, NAV-2729 reduced SARS-CoV-2 viral loads also in more physiologic models of infection: Calu-3 and kidney organoids. This highlighted the importance of ARF6 in multiple cell contexts. Together, these experiments points to ARF6 as a putative target to develop antiviral strategies against SARS-CoV-2.

## Introduction

SARS-CoV-2 is a newly emerged beta-coronavirus and causative agent of the COVID-19 pandemic. While several vaccines and antivirals have been approved to limit severe COVID-19, the spread of the virus is not yet under control. This highlights the importance of developing additional therapeutic strategies that can complement vaccine rollout, especially at the epidemic centers. Given the propensity of RNA viruses to develop drug resistance, there is an unmet need for additional antiviral compounds that can curb the infection and transmission by targeting additional steps in the virus life cycle. Viral entry is a step in the viral cycle that is promising for antiviral development. This important step of the viral life cycle has been characterized very early after the isolation of SARS-CoV-2, leading to the discovery of the uncoating receptor, ACE2 [1] and the two entry pathways [2]: via plasma membrane fusion or endocytosis. Membrane fusion at the plasma membrane occurs after SARS-CoV-2 surface (S) protein activation by proteinase cleavage, including TMPRRS2, whereas fusion in the endocytic compartment requires the intracellular protease cathepsin L. The pathway and molecular players that govern endocytosis-mediated entry are however poorly characterized. Endocytosis of viruses can occur via different pathways, including clathrin-mediated endocytosis (CME), caveolar or lipid raft-mediated endocytosis, macropinocytosis and variations of these themes [3]. Some viruses are able to use more than one pathway, which introduces a further level of ambiguity to the classification of virus entry pathways. Research on SARS-CoV-2 endocytic-mediated entry has been scant. A single report using a lentivirus system in ACE-2 overexpressing 293T cells suggests that SARS-CoV-2 endocytosis is dependent on clathrin and dynamin [4]. However, studies on SARS-CoV-2 endocytic entry pathway in the context of a productive infection were missing at the beginning of our investigation.

ADP-ribosylation factor 6 (ARF6) is a multi-functional cellular protein. It directly activates lipid modifying enzymes, such as phosphatidylinositol 4-phosphate 5-kinase, and it stimulates actin polymerization. ARF6-positive cargoes can be directly transported on microtubule filaments; and it also assists autophagy, particularly the initiation and autophagosome formation [5]. Because of its roles in endocytosis and trafficking, ARF6 is co-opted by several pathogens; bacteria (e.g., enteropathogenic and enterohemorrhagic *Escherichia coli, Salmonella Typhimurium, Shigella*) and viruses (e.g., HIV-1, Coxsackievirus A and B, Epstein Barr virus). ARF6 was recently identified as a cellular interaction partner of the uridine-specific endoribonuclease Nsp 15 of SARS-CoV-2 [6] and a recent re-analysis of a RNA sequencing dataset of infected A549-ACE2 cells, ARF6 is involved in three of the four identified modules mediating host cell responses during SARS-CoV-2 infection, i.e., viral entry, regulation and signaling, and immune responses [7]. We therefore sought out to determine the role of ARF6 in SARS-CoV-2 endocytosis and, more broadly, infection.

We first found that in the human hepatocarcinoma cell line, Huh-7 SARS-CoV-2 is resistant to camostat, a specific TMPRSS2 inhibitor, and hence SARS-CoV-2 entry mostly relies on endocytosis for infection. By genetic depletion and pharmacologic inhibition, we further demonstrated that SARS-CoV-2 entry into Huh-7 is dynamin-independent but dependent on cholesterol and ARF6. This host factor is also important for infection in the more physiologic relevant models of human lung Calu3 cells and kidney organoids, suggesting that ARF6 might be an effective therapeutic target.

## Material and Methods

### Cells and virus

Human hepatocarcinoma cells, Huh-7, and human lung adenocarcinoma cells, and Calu3, were obtained by collaborators and maintained at 37 °C and 5% CO2 in Dulbecco’s modified Eagle’s medium (DMEM) (Gibco), supplemented with 10% heat-inactivated fetal bovine serum (FBS), Hepes, nonessential amino acids, L-glutamine, and 1× Pen-Strep (Gibco). Kidney organoids were generated using human embryonic stem cells (UM77-2), as previously described [8]. SARS-CoV-2 was obtained through BEI Resources (SARS-Related Coronavirus 2, Isolate USA-WA1/2020, NR-52281 and Isolate hCoV-19/USA/GA-EHC-2811C/2021 [Omicron Variant] NR-56481). Lack of genetic drift of our viral stock was confirmed by deep sequencing. Viral titers were determined by TCID_50_ assays in Vero E6 cells (Reed and Muench method). All experiments using SARS-CoV-2 were performed at the University of Michigan under Biosafety Level 3 (BSL3) protocols in compliance with containment procedures in laboratories approved for use by the University of Michigan Institutional Biosafety Committee and Environment, Health and Safety.

### SARS-CoV-2 infection and quantification by immunofluorescence

Cells were seeded at 3,000 cells/well (384-well plate, Perkin-Elmer, 6057300) or 10,000 cells/well (96-well plates, Costar, 3916) and allowed to adhere overnight. Compounds were then added to the cells and incubated for 2 hrs at 37 °C. Plates were then transferred to BSL3 containment and infected with SARS-CoV-2 WA1 or Omicron at a MOI of 1. In the case of infection with the Omicron variant, a pre-treatment of the virus with 10 μM of porcine trypsin (Millipore-Sigma, cat nr. T0303-1G) for 15 min at 37’C was included. After 1 h of absorption on cell (final concentration of trypsin was 2 μM), the virus inoculum was removed and fresh media with inhibitors was added. Uninfected and vehicle-treated infected cells were included as negative and positive controls, respectively. Plates were then stained as previously described [9] by using anti-nucleocapsid protein (anti-N) SARS-CoV-2 antibody (Antibodies Online; catalog no. ABIN6952432) followed by staining with secondary antibody Alexa-647 (goat anti-mouse; Thermo Fisher; A21235) and DAPI for nuclei staining. Stained cells were acquired on Thermo Fisher CX5 high-content microscopes with a 20×/0.45 N.A. LUCPlan FLN objective. Laser autofocus was performed and 9 or 18 fields per well were imaged covering ~80% of the well area. Images were analyzed with Cell Profiler software and percent of infected cells (N-positive) was calculated on the total cell count (i.e., DAPI positive).

### SARS-CoV-2 infection and quantification by TCID_50_ assay

200,000 Huh-7 or Huh-7 knock-out (KO) cells were seeded in 12-well plates. For the Calu-3, 100,000 cells were seeded in 24-well plates. The next day or at 80% cell confluency, compounds were added to the cells and incubated for 2 hrs. Cells were then infected with SARS-CoV-2 WA1 at a MOI of 1 for 1 h at 37 °C. Inoculum was removed, cells were washed 3 times with PBS without Magnesium and Calcium (PBS-/-) and fresh medium was added. Cells were harvested 1 day post infection (dpi) and lysates were obtained by one cycle of freeze-thaw. For the kidney organoids, 3D-spheres were collected in 1.5 mL tubes and infected with SARS-CoV-2 at an estimated MOI of 1. After infection, 3D-spheroids were washed 3 times with PBS-/- and gentle centrifugation (80 g for 2 min). A minimum of four 3D-spheroids were transferred in a 96-well plate and medium with selected compounds was added to the cells. SARS-CoV-2 infection was quantified by TCID_50_ on pre-seeded Vero E6 cells (Reed and Muench method) or by RT-qPCR (IDT technologies, Charité kit) after RNA extraction with TRI reagent and Direct-Zol RNA miniprep kit (Zymo Research).

### Generation of CRISPR-KO cell lines

Individual guide RNA (gRNA) sequences (**Supplemental Table 1**) were cloned into BsmBI-digested lentiCRISPRv2 (Addgene #52961, a gift from Feng Zhang [10]) and resulting lentiviral stocks prepared as previously described [11]. Huh-7 and Calu-3 cells were transduced with lentiviral stocks at an MOI of ~1, treated 1 day later with puromycin (3 ug/mL) until death of control uninfected cells was complete, and passaged as needed to maintain logarithmic phase growth for 2-3 weeks. Cells were then either harvested for mRNA and protein preparations or infected with SARS-CoV-2 infection as described above.

### Attachment/internalization assay

Huh-7 or ARF6-KO cells were plated in 48-well plates at 100,000 cells per well and allowed to adhere overnight. The following day, NAV-2729 was added at the indicated concentration and incubated for 2 hrs. Following compound incubation, cells were infected with SARS-CoV-2 at an MOI of 10 for 1 h at 4 °C to allow for viral binding. Cells were then washed three times with ice-cold PBS-/- to remove unbound virus and harvested in TRI reagent. Another set of treated-infected cells was transferred at 37°C to allow for viral internalization. One hour after incubation, cells were treated with trypsin to remove bound-not internalized virus and after centrifugation at 500 x g for 5 min, cells were harvested in TRI reagent. RNA was extracted by using the Direct-Zol RNA miniprep kit (Zymogen; R2052) and viral RNA was quantified by RT-qPCR. Proportion of internalized virus was calculated over the bound virus.

## Results

### SARS-CoV-2 infection in Huh-7 is resistant to camostat treatment but sensitive to Z-FA-FMK

To investigate SARS-CoV-2 endocytosis, we screened a range of cell lines for the antiviral activity of camostat mesylate, a specific inhibitor of TMPRSS2 (data not shown). Infection readout was carried out with an imaging-based pipeline by using a SARS-CoV-2 anti-nucleocapsid (N) antibody to detect infected cells. Interestingly, in the human hepatocarcinoma cell line Huh-7, SARS-CoV-2 was completely resistant to the antiviral action of camostat (**Figure 1A**). In contrast, treatment with the irreversible cysteine protease inhibitor Z-FA-FMK, an inhibitor of the intracellular protease cathepsin L, resulted in a strong antiviral activity with EC50 of 11 nM (**Figure 1B**) [9]. Representative images of SARS-CoV-2-infected Huh-7 in the presence of Z-FA-FMK and camostat are available in **Figure 1C**. These data suggest that SARS-CoV-2 entry into Huh-7 cells is likely mediated by endocytosis and not fusion at the plasma membrane. Therefore, Huh-7 cells represent a model cell line to mechanistically study SARS-CoV-2 endocytic entry.

**Figure 1:**
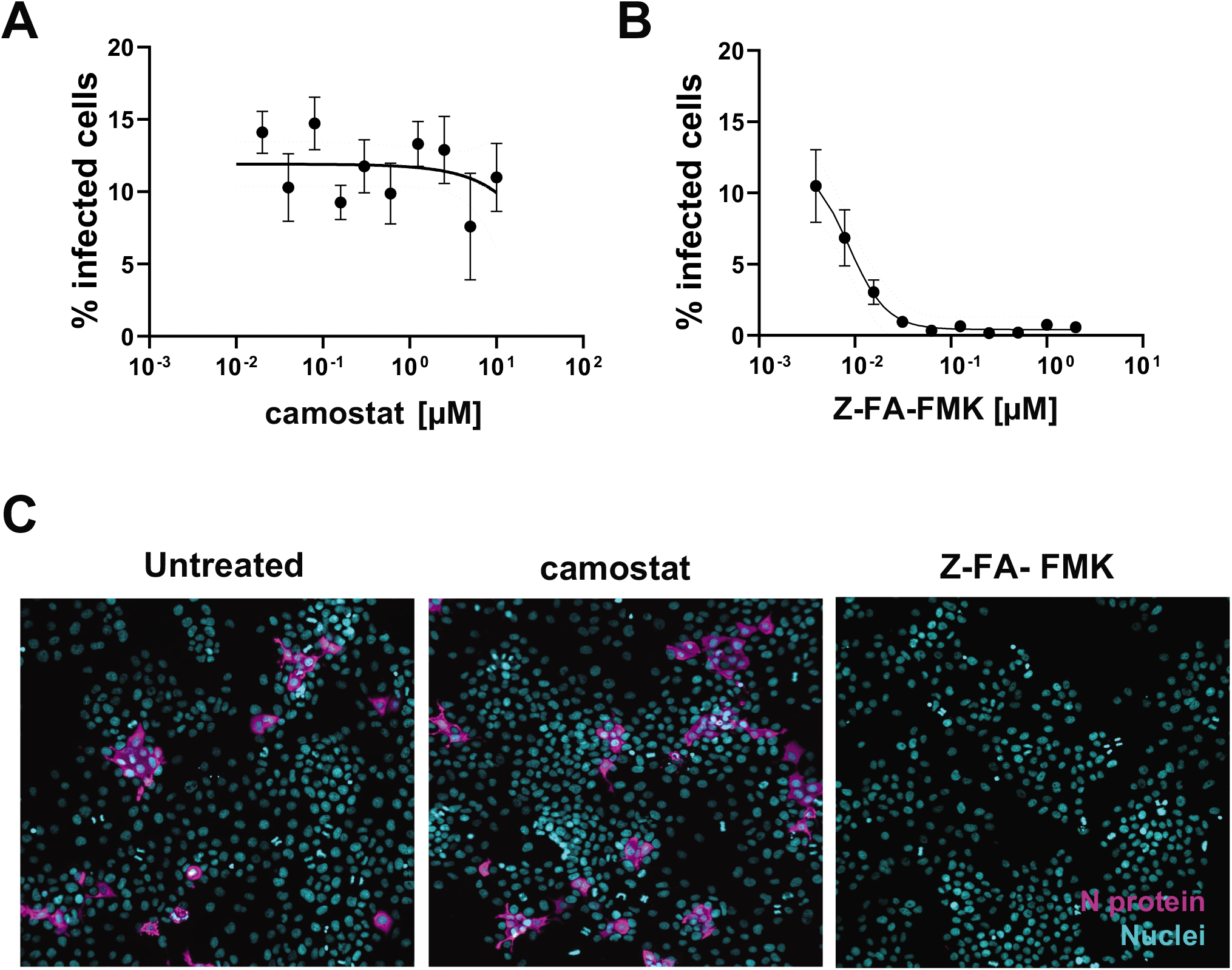
Huh-7 cells are resistant to the antiviral effect of camostat mesylate. A-B) Dose-response curves of camostat mesylate (A) and the cathepsin B inhibitor Z-FA-FMK (B). Huh-7 cells were pre-treated with compound and infected with SARS-CoV-2 at MOI of 1 for 1h at 37°C. 48 hpi cells were fixed and permeabilized and stained with anti-N antibody. Cells were imaged with high content imaging platform CX5 and analyzed with Cell profiler for image segmentation. The % of infected cells were normalized to the non-treated infected condition. Graph represents average SD of N=3 independent experiments with n=3 technical replicate, each. C) Representative images of Huh-7 cells infected with SARS-CoV-2 or infected/treated with camostat (500 nM) and Z-FA-FMK (500 nM). N protein staining is represented in magenta, nuclei are in blue.

### SARS-CoV-2 uptake into Huh-7 cells is dynamin-independent

A previous report with a lentivirus system in ACE-2 overexpressing 293T cells, suggests that SARS-CoV-2 endocytosis is dependent on clathrin and dynamin [4]. We therefore determined if dynamin governs SARS-CoV-2 entry into Huh-7 cells by using dynasore, a specific GTPase inhibitor of dynamin I, II and Drp1 (the mitochondrial dynamin) [12]. ß-methyl cyclodextrin, a cholesterol-removing agent that mostly blocks formation of lipid rafts important for endocytosis, was used as a positive control since ACE2 is present in lipid rafts [13] and SARS-CoV entry is dependent on cholesterol [14]. Huh-7 cells were pretreated 2 hrs prior to infection with selected compounds at non-toxic concentrations (**Supplementary Figure 1**) and infection with SARS-CoV-2 was quantified by N protein expression. As expected, treatment with ß-methyl cyclodextrin blocked viral infection. However, in contrast with previous findings, treatment with the specific dynamin inhibitor dynasore, revealed a significant increase in the percentage of infected cells (**Figure 2)**. This observation points to a dynamin-independent but cholesterol-dependent endocytic uptake mechanism of SARS-CoV-2 into Huh-7 cells.

**Figure 2:**
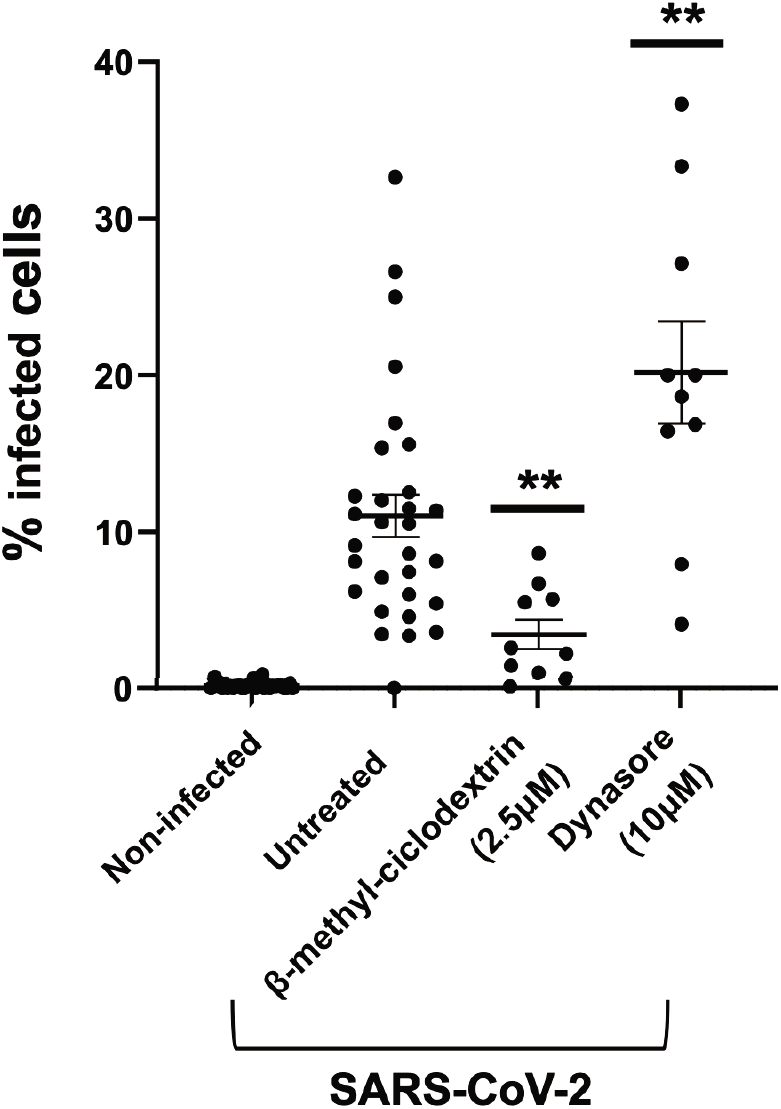
Screening of inhibitors of the dynamin-dependent entry pathways in Huh-7 cells. Huh-7 cells were pre-treated with compounds at the indicated concentration and infected with SARS-CoV-2 at MOI of 1 for 1 h at 37°C. 24 hpi cells were fixed, permeabilized and stained with anti-N antibody. Cells were imaged with high content imaging platform CX5 and analyzed with Cell profiler for image segmentation.

### Genetic perturbation of ARF6 results in a reduction in the percentage of infected cells

Dynamin-independent pathways are governed by the Rho GTPase-activating protein 26 (GRAF-1), flotillin (FLOT-1) or ADP-ribosylation factor 6 (ARF6) [15]. Since ARF6 was recently described for mediating host cell responses during SARS-CoV-2 infection, we generated two polyclonal populations of CRISPR-targeted cells with gRNAs specific for ARF6 (clones 1 and 2). ARF6 depletion was confirmed by RT-qPCR, whereby the efficacy of knock-out (KO) was measured as a percentage of mRNA target gene expression compared to the non-targeting guide (NTg) Huh-7 cell line (for clone 1, c1=39% and clone 2, c2=56%) and by Western blot (**Supplementary Figure 2A**). To test the efficacy of the CRISPR pipeline, an ACE2-KO Huh-7 line was generated in parallel as a control. Cells were infected with SARS-CoV-2 at MOI of 1 and harvested 1 and 2 dpi. A combination of TCID_50_ assay and RT-qPCR were used to quantify viral infection and replication, respectively (**Figure 3A and 3B**). No significant differences were detected by TCID_50_ between ARF6 KO and NTg cells whereas replication was ablated in the control ACE2-KO Huh-7 cells. To independently validate these findings, an imaging-based readout for SARS-CoV-2 infection was used, since it enables analysis at the single cell level. The assay was also adjusted to a 1 dpi readout and infecting cells with SARS-CoV-2 at the higher MOI of 10. For ease of experimentation, only ARF6-KO c1, with lower expression of the target gene was used for this analysis. Representative images at 1 dpi are shown in **Figure 3C**. Intriguingly, we observed a significant decrease in the percentage of infected cells in ARF6 KO than the NTg cells, suggesting that ARF6 might be involved in SARS-CoV-2 entry (**Figure 3C**). Collectively, these data suggest a role for ARF6 in SARS-CoV-2 infection in Huh-7.

**Figure 3:**
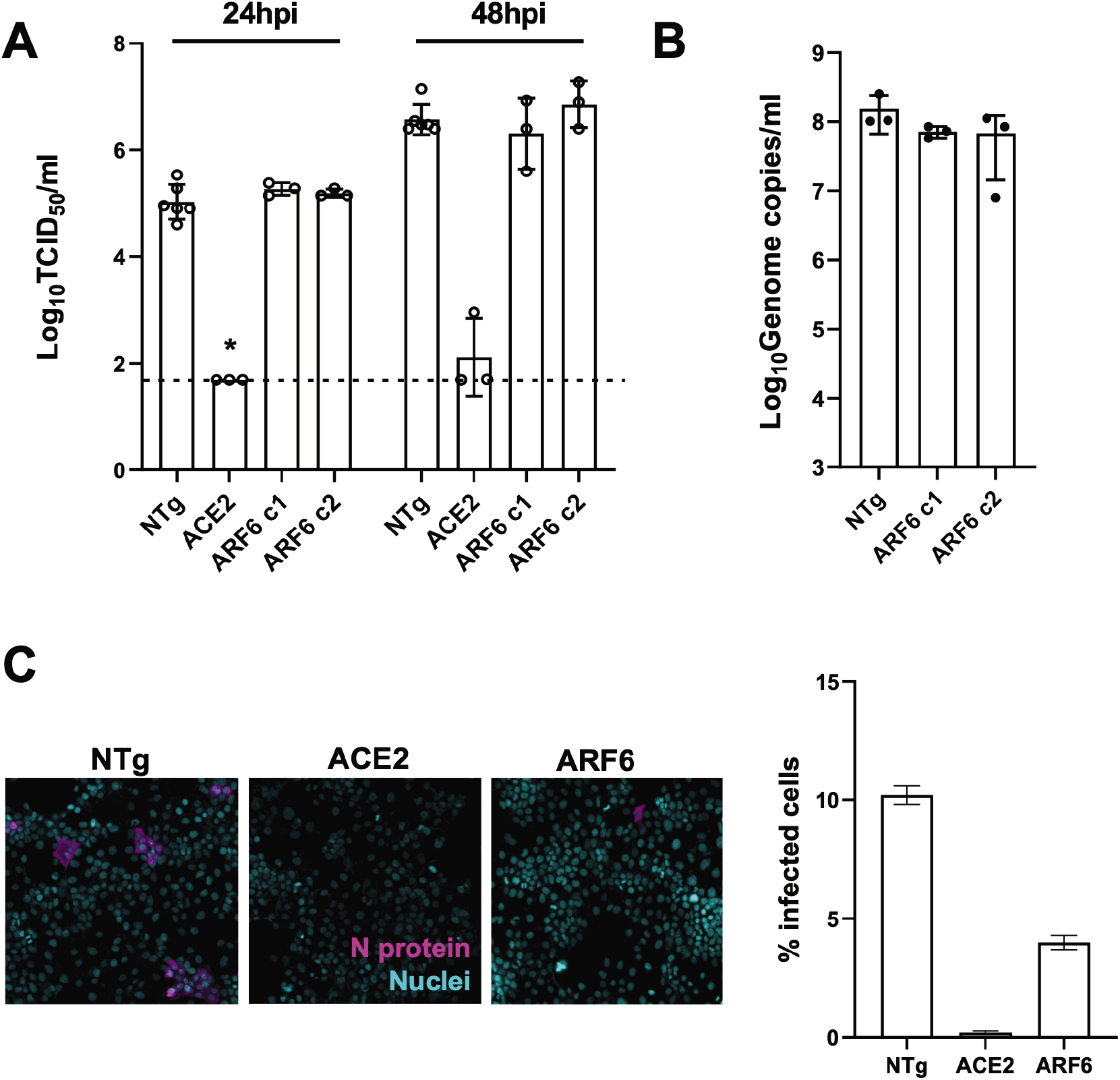
Genetic depletion of ARF6 does not affect viral titers but reduce the percentage of infected cells A-B) Viral titer determination of SARS-CoV-2-infected NTg, ACE2-KO ARF6-KO Huh7 clones (MOI of 1) by TCID_50_/ml at 24 and 48 hpi (A) and by RT-qPCR at 24dpi (B). ACE2-KO was used as a negative control of infection. Viral infectious titer was determined by TCID_50_ on Vero E6 cells. Dashed line represents the limit of detection of the TCID_50_ assay. C) Representative images of SARS-CoV-2-infected NTg and KO-Huh7 clones (MOI of 10). N protein staining is represented in magenta, nuclei are in blue (DAPI). The graph represents the quantification of SARS-CoV-2 infected cells (MOI of 10) at 24 hpi. Cells were imaged with high content imaging platform CX5 and analyzed with Cell profiler for image segmentation. The % of infected cells was normalized to the non-treated infected condition. Graph represents average SD of N=3 independent experiments with n=3 technical replicate, each.

### Pharmacologic inhibition of ARF6 blocks SARS-CoV-2 internalization

ARF6 has been the object of extensive studies and specific small molecule inhibitors are available commercially. We therefore selected the inhibitor NAV-2729 to test its effect on SARS-CoV-2 infection. Huh-7 cells were pre-treated with a 1:2 dilution series of compounds and infected with SARS-CoV-2 at MOI of 1. NAV-2729 showed a dose-dependent inhibition of infection (**Figure 4A**), suggesting that ARF6 is an important factor for SARS-CoV-2 infection in Huh-7. We next selected an efficacious non-toxic concentration of NAV-2729 (**Supplementary figure 1A**) to confirm inhibition by TCID_50_ (**Figure 4B**). Treatment with NAV-2729 resulted in a ~2.5-Log reduction in viral infection in the NTg control. We also wanted to assess whether viral entry into ARF6-KO switched to a dynamin-dependent entry mechanisms but treatment of the ARF6-KO cells with dynasore did not result in changes to infection, further corroborating that dynamin is not involved in SARS-CoV-2 endocytosis in Huh-7 (**Figure 4B**). To determine whether the NAV-2729 inhibitor worked at the level of viral internalization, an attachment/internalization assay was performed. Briefly, NTg cells were pre-treated with NAV-2729 and infected with SARS-CoV-2 at the MOI of 10. ARF6-KO cells were included as a control. One hour post-inoculation on ice, cells were washed and one set was harvested to quantify the virus attached at the cell surface by RT-qPCR. While treatment with NAV-2729 did not affect virus binding to NTg cells, ARF6-KO cells showed a significantly higher level of SARS-CoV-2 binding compared to NTg cells (**Figure 4C**). The other batch was incubated at 37°C for 1 hr to allow for viral internalization, and after bound-not internalized virus was removed by a treatment with trypsin, cells were harvested to quantify the internalized virus by RT-qPCR. To control for differences in SARS-CoV-2 binding between cells, we calculated the ratio of internalized *versus* bound virus (**Figure 4D**). NAV-2729 reduced viral internalization by 75% *versus* the more modest 25% observed in the ARF6-KO, suggesting that ARF6 is a factor that governs endocytosis in Huh-7.

**Figure 4.**
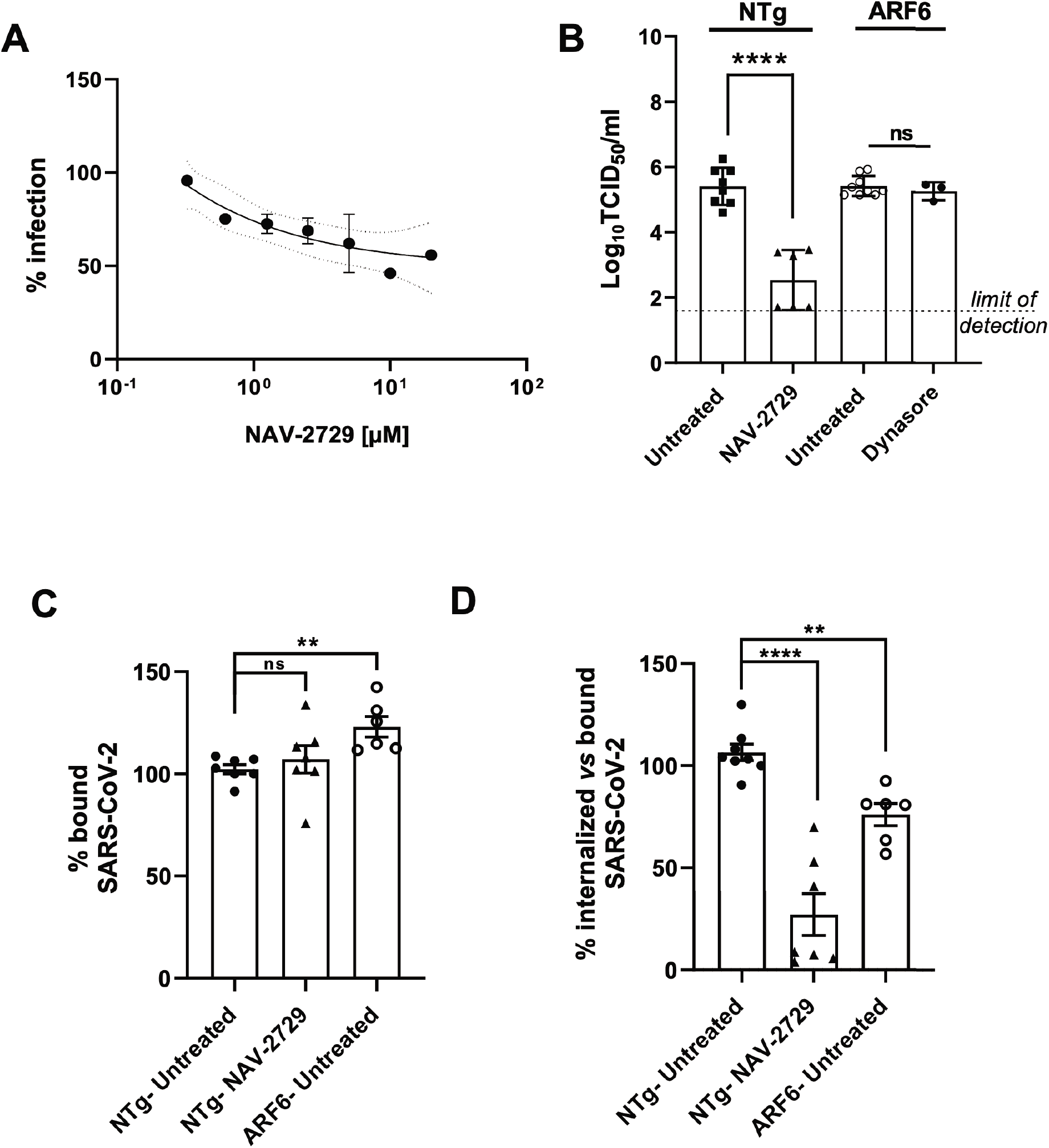
ARF6 is important during SARS-CoV-2 internalization in Huh-7 cells. A) NAV-2729, an ARF6 inhibitor was used to pre-treat Huh-7 cells in a 384-well plate 2 hrs prior infection with SARS-CoV-2 (MOI of 1). Cells were harvested 48 hpi and subjected to the imaging pipeline for viral quantification. 1:2 dose response curves were performed in triplicate with 3 technical replicates, each. B) NTg Huh-7 were pre-treated with NAV-2927 (10 μM) and ARF6 KO were pre-treated with dynasore (10 μM) for 2 h before infection with SARS-CoV-2 (MOI of 1). Virus inoculum was removed 1 hpi and cells were incubated with fresh medium +/− compounds. Cells were harvested 24 hpi and viral titer was determined by TCID_50_. C-D) NAV-2729 was used to pre-treat Huh7 cells 2 hrs prior infection with SARS-CoV-2 (MOI of 10) on ice. NTg and ARF6-KO Huh7 cells were used as a control. 1 h post infection on ice, cells were washed twice and one set was harvested in TRI reagent. The second set was incubated at 37’C to allow viral internalization. One hour post incubation, cells were treated with trypsin to remove bound, non-internalized virus and were harvested in TRI reagent. Bound and internalized virus were determined by RT-qPCR: Graph in panel C shows % of bound virus over untreated control (NTg Huh-7), whereas graph in panel D measures the internalized fraction of SARS-CoV-2 over the bound fraction. Statistical t-test was performed on GraphPad Prism on N= 3 biological replicate with at least 2 technical replicates, each. **p-value ≤0.01, **** p-value ≤0.001

### ARF6 is an important host factor for SARS-CoV-2 infection in more physiological relevant infection models

To test the importance of ARF6 in other cellular contexts, we used the ARF6 inhibitor NAV-2729 in more physiologic relevant models that are sensitive to camostat: human renal organoids and human lung Calu-3 cells [16]. In both models, treatment with non-toxic concentration of NAV-2729 (**Supplementary figure 1B**) resulted in a significant reduction of viral replication (**Figure 5A and 5B**). The nucleoside inhibitor remdesivir was used as a control. In addition, we performed CRISPR-Cas9 knockout of ARF6 in Calu-3 cells (**Supplementary figure 2B**) to corroborate the pharmacologic studies with NAV-2729. Infection with SARS-CoV-2 at an MOI of 1 was less efficient in the Calu3 line transduced with the ARF6-targeting guide (ARF6) than in Calu-3 cells transduced with a non-targeting RNA guide (NTg) (**Figure 5B**). Although the data did not reach statistical significance, the decrease in SARS-CoV-2 infection in Calu3 ARF6-KO cells was more pronounced than in Huh-7 ARF6 KO cells (compare **Figure 5B** with **Figure 3A**), suggesting that ARF6 might play additional roles in Calu-3 cells compared to Huh-7.

**Figure 5.**
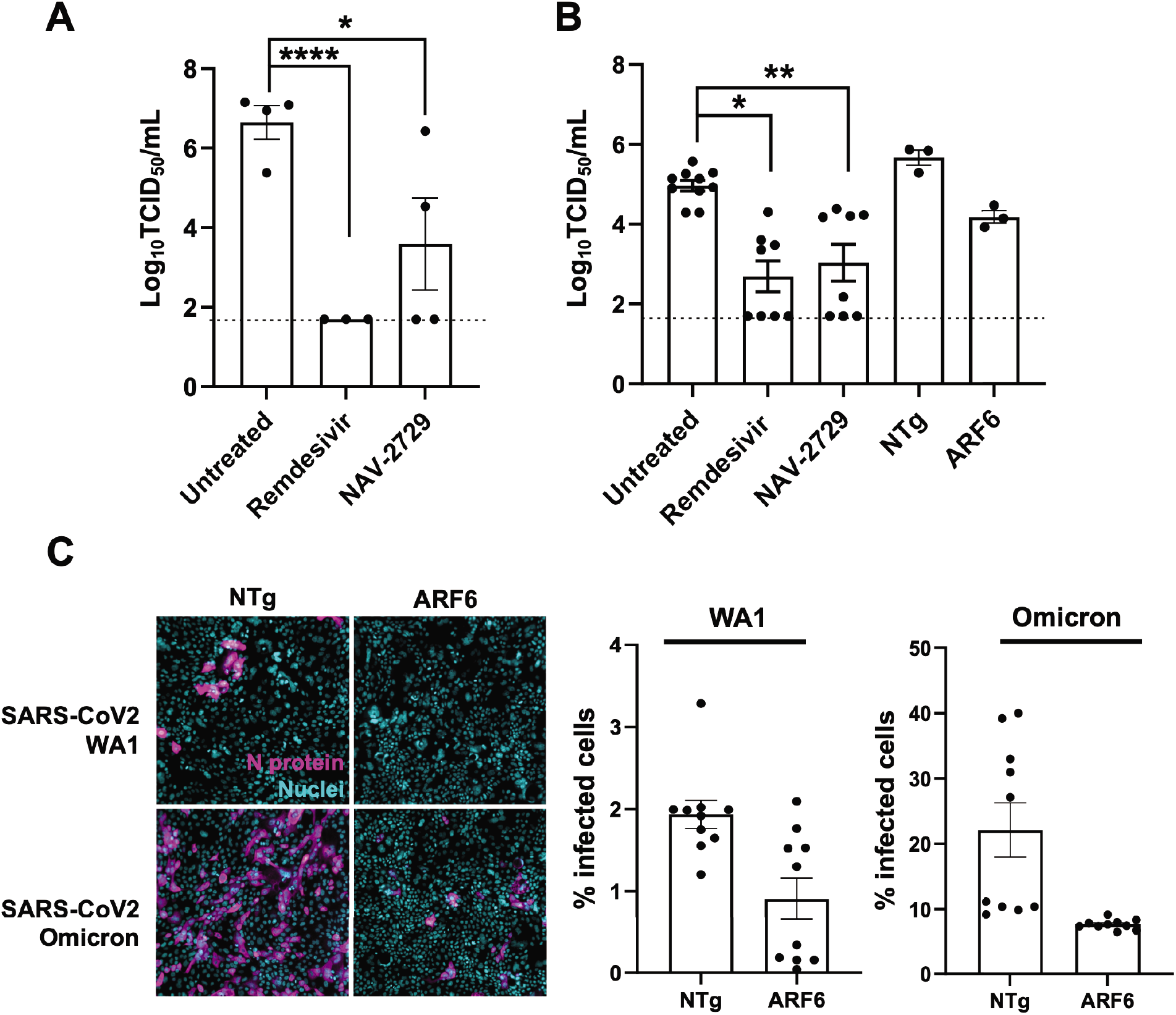
ARF6 is critical for SARS-CoV-2 infection in more physiologic relevant cell types. A) Human kidney organoids were collected and resuspended in maintenance media. Groups of four organoids were infected with SARS-CoV-2 at MOI of 1. Organoids were washed twice and put in culture on a matrigel-coated plate for 48 hrs. Cells were then harvested, and virus was quantified by TCID_50_. Each dot represents a technical replicate of N=2 biologically independent experiments. B) Calu-3 cells and Calu-3 transduced with a non-targeting guide (NTg) or ARF6-targeting guide (ARF6) were seeded in 48-well plate and allowed to reach 80% confluence. SARS-CoV-2 WA1 was used to infect cells at MOI of 1 for 24 hrs. Cells were then harvested, and virus was quantified by TCID_50_. Each dot represents a technical replicate. C) Representative images (left) and quantification (right) of Calu3 transduced with a non-targeting guide (NTg) or ARF6-targeting guide (ARF6) and infected with trypsin activated SARS-CoV-2 WA1 or SARS-CoV-2 Omicron (MOI of 1) for 24 hrs. Cells were fixed, permeabilized and stained with anti-N primary antibody and anti-mouse AF647 secondary antibody (N protein, magenta) and Hoechst (nuclei, blue). Cells were imaged with the high content imaging platform CX5 and analyzed with Cell profiler for image segmentation. Each dot represents the average of % of infected cells over 20 fields of image per well. Each dot represents a technical replicate.

Lastly, to shed more light on the mechanism by which ARF6 assists SARS-CoV-2 replication in Calu-3 cells, we infected NTg and ARF6 KO Calu-3 cells with SARS-CoV-2 Omicron variant. This variant has been demonstrated to rely primarily on endocytosis and not on membrane fusion for entry *in vitro* and *in vivo* [17]. Both virus isolates were pre-treated with trypsin prior to infection to proteolytically cleave the S protein and bypass the need of cellular proteases [18]. Under these conditions, infection of NTg Calu-3 cells with SARS-CoV-2 Omicron resulted in ~15-fold higher infection (as measured by % of N-positive cells at 24 hpt) than with the WA1 strain (**Figure 5C**). Infection of ARF6 KO with both strains showed a significant reduction compared to NTg cells, suggesting that ARF6 might play a role in post-entry steps in Calu-3 cells. Overall, these results point to the importance of ARF6 in SARS-CoV-2 infection and to a putative therapeutic target against coronavirus infection.

## Discussion

In this study, we uncovered that ARF6 is a critical player of SARS-CoV-2 infection in cells of liver, lung, and kidney origin. In addition, in Huh7 cells, ARF6 inhibited SARS-CoV-2 uptake with a mechanism that is further dependent on cholesterol but independent of dynamin and by extension likely clathrin-independent. This is consistent with a prior study that excludes the involvement of dynamin, clathrin, caveolin and endophilin A2 for entry of SARS-CoV-2[14] and with the entry pathway identified for SARS-CoV, which is cholesterol-dependent but clathrin- and caveolin-independent [19]. However, our finding differs from a study demonstrating the importance of clathrin in SARS-CoV-2 entry [4]. That study was performed with a lentivirus system in 293T cells and not in the context of authentic SARS-CoV-2 infection. Thus, differences in virus and/or cell type may alter the endocytic uptake pathway. During the revision of this manuscript, Zhou et al. also published that ARF6 is an important host factor for SARS-CoV-2 entry [20]. Specifically, their study used an siRNA approach and SARS-CoV-2 pseudovirus to demonstrate a 2-fold reduction of viral entry in Huh-7 cells. This is consistent with the levels of reduction that we observe in the context of SARS-CoV-2 infection in ARF6 KO cells (by immunofluorescence) and upon pharmacological inhibition of ARF6 by the specific inhibitor NAV-2729. We posit that the observed effect of ARF6 on infectivity in Huh-7 cells by immunofluorescence but not in infectious virus titers in Figure 3 might be simply due to the tenfold higher MOI used in the former experiment given the heterogeneity of the knockdown. Intriguingly, we observed that CRISPR-depleted cells exhibited a less pronounced phenotype as compared with those treated with the ARF6 specific inhibitor, NAV-2729. This could be due to several reasons. Because of the redundancy of endocytic pathways and lack of an important intracellular trafficking factor, prolonged passaging of the ARF6-KO cells and the potential incomplete activity of the guide RNA as indicated by the qPCR data might have led to compensatory mechanisms and a less uniform phenotype. In contrast, the short-term treatment with a small molecule inhibitor at non-toxic concentrations does not affect or modify the physiology of the cell and reduces the impact of compensatory mechanisms and could thus be more relevant for the study of these redundant endocytic pathways.

In our investigation, we expanded on the importance of ARF6 also in the context of infection of other cell types; kidney organoids and Calu3 cells, a model of respiratory epithelial cells. In both systems, the ARF6 inhibitor, NAV-2729, showed a strong antiviral effect. These cell models support SARS-CoV-2 entry also via membrane fusion [21]. ARF6-KO Calu3 or Huh7 cells exhibited a reduction of infection by both immunofluorescence and TCID_50_. These data highlight the important role for ARF6 during SARS-CoV-2 infection. Considering the multiple functions of ARF6 in intracellular membrane trafficking [5] and the observation that the decrease in SARS-CoV-2 infection in Calu3 ARF6-KO cells was more pronounced than in Huh-7 ARF6 KO cells (compare Figure 5B with Figure 3A), we cannot rule out a role for ARF6 in post-entry steps of the viral life cycle. However, additional studies are needed in the future to examine this possibility.

It is not known what proportion of SARS-CoV-2 virions is entering via membrane fusion at the plasma membrane or endocytosis *in vivo*. SARS-CoV-2 prefers activation by TMPRSS2 but if the target cells express insufficient TMPRSS2 or if a virus–ACE2 complex does not encounter TMPRSS2, ACE2-bound virus is internalized via endocytosis [2]. TMPRSS2 is found in the gastrointestinal, respiratory, and urogenital epithelium but not in other extra-mucosal compartments (i.e., heart, liver, kidneys, brain, spleen, and lymph nodes), where virus RNA or antigen is found in severe and deadly cases of COVID-19, suggesting that entry and replication of SARS-CoV-2 in these organs might be exclusively driven by endocytosis. It is also possible that SARS-CoV-2 entry occurs by plasma membrane fusion and endocytosis, regardless of the compartment. Therefore, it would be of interest to use NAV-2729 in a small animal model to investigate the *in vivo* relevance of the endocytic pathway and its importance in driving severe disease. Altogether, these data highlight ARF6 as a potential target in the development of therapeutics against COVID-19. Future studies focusing on investigational drugs targeting ARF6, such as Linsitinib that has antiviral activity in a pseudovirus assay [7], could hold promise for the development of additional SARS-CoV-2 therapeutics.

## Acknowledgement and funding information

We thank Dr. A. Lauring (University of Michigan) for deep sequencing of the viral stock. The work was in part funded by the University of Michigan Biological Science Scholars Program to C.E.W. C.M. was supported by the Marie-Skłodowska-Curie Actions action global fellowship (GA 841247). JZS is supported by the National Institute of Diabetes and Kidney Diseases (R01DK120623) and National Center for Advancing Translational Sciences Grant UL1TR002240. J.W.W. is supported by the Pharmacological Sciences Training Program T32 Training Grant GM007767.

## Conflict of Interest

The authors declare that there are no conflicts of interest.

## Figure legends

**Supplementary figure 1:**
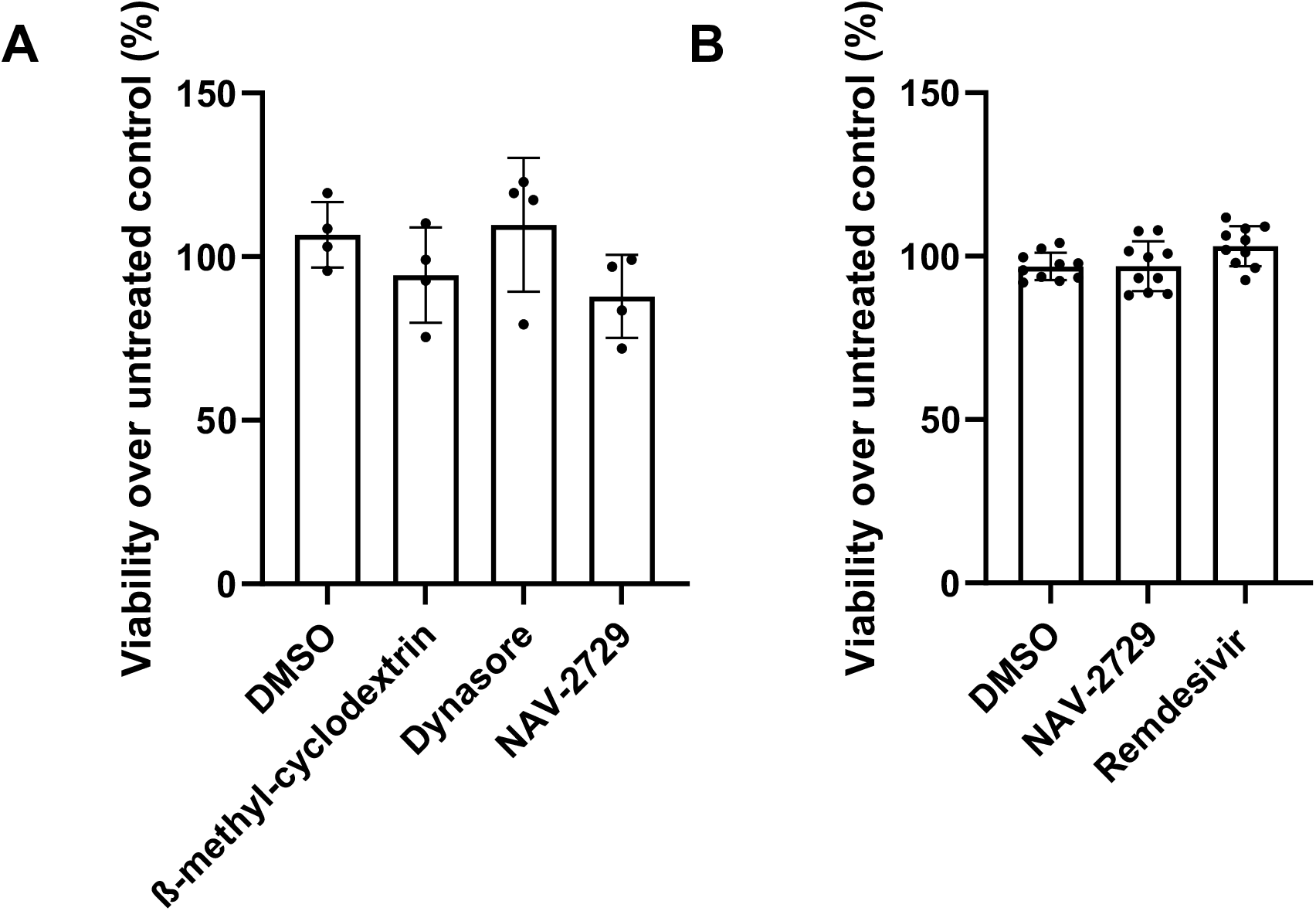
A-B) Toxicity assay of selected drugs by Resazurin assay in Huh-7 (panel A) and Calu-3 cells (panel B).

**Supplementary figure 2:**
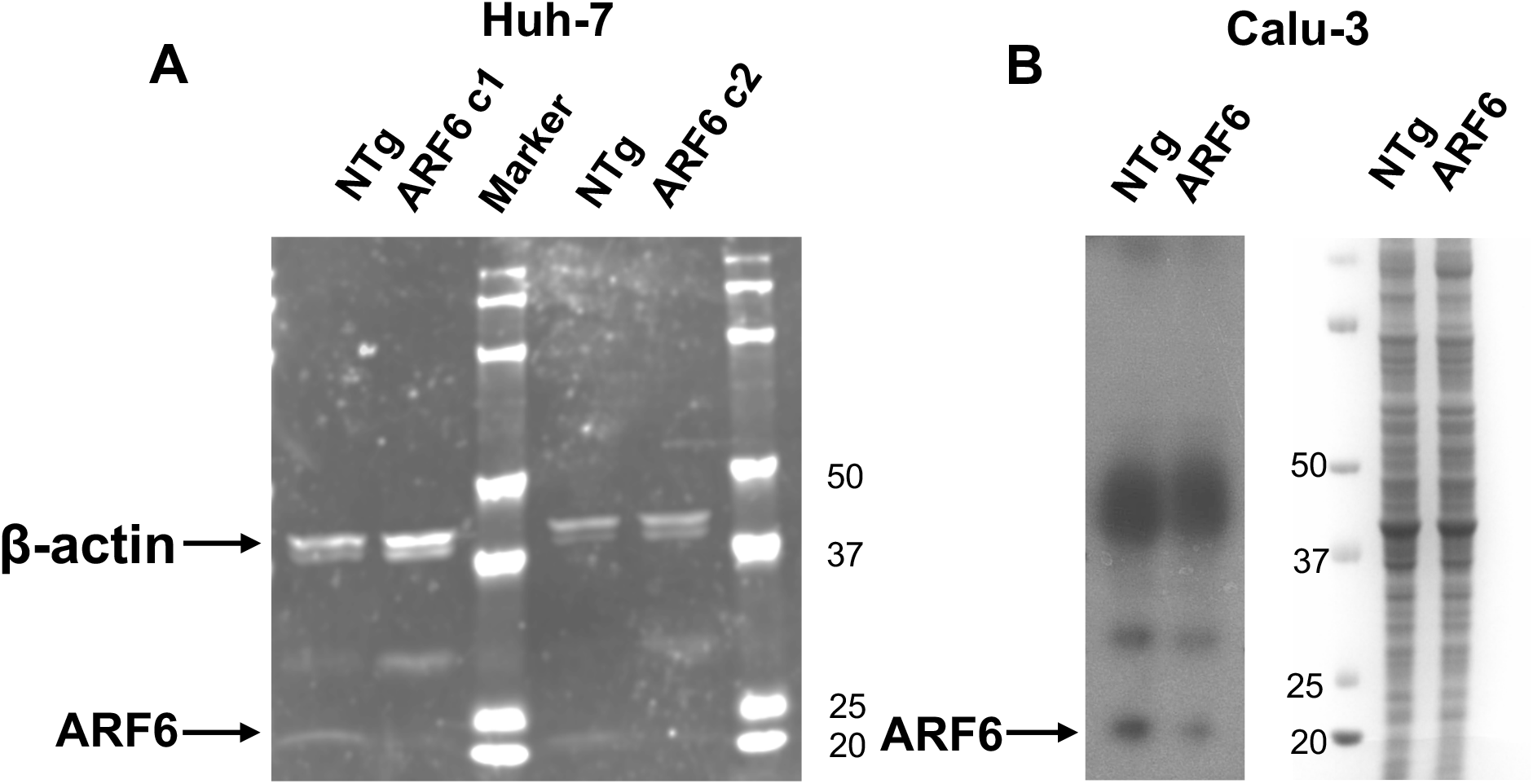
Western blot for ARF6 expression. A) Huh-7 lysates from non-targeting (NTg) and 2 clones of ARF6-knock-out (KO) preparations were obtained from cells seeded overnight in a 6-well plate and analyzed for ARF6 expression by Western blot. ß-actin was used as a loading control. Molecular weight marker (in kDa) is shown on the right. B) NTg and ARF6-KO Calu-3 cells were similarly obtained and analyzed by Western blot. As a loading control, a Coomassie blue staining of the gel was performed. Molecular weight marker (in kDa) is shown in the middle.

**Supplementary Table 1:**
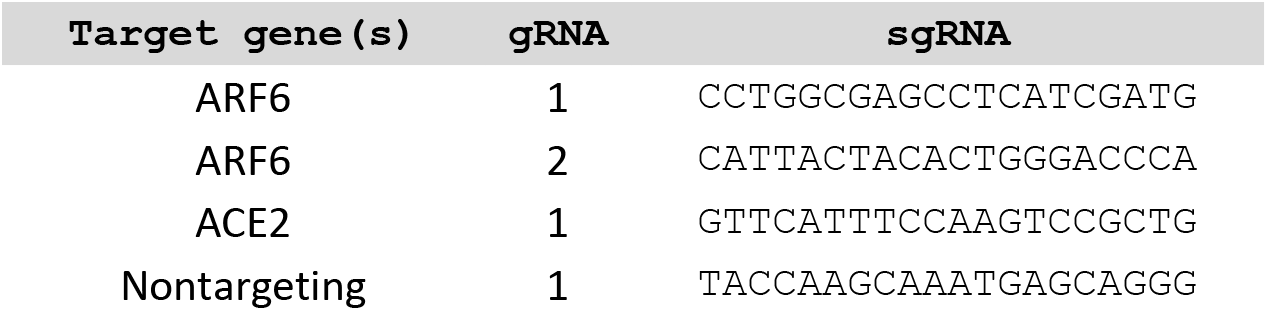
Sequences of sgRNAs for targeted CRISPR/Cas9 knockout.

## Supplementary Methods

### Western blot

Cells were lysed in RIPA buffer (Pierce, 89900) with PhosSTOP (Roche, 04906845001) and cOmplete EDTA-free protease inhibitor cocktail (Roche, 11873580001) on ice. After vortexing to ensure a complete lysis, cells were centrifuged at 4°C, 14,000 × g for 15 min. The supernatant was mixed with β-mercaptoethanol-containing (10%) Laemmli buffer (Bio-Rad, cat nr. 1610747) at 3:1 ratio and denatured at 90°C for 10 min with a heat block. Samples were spun down before being loaded in a 4 to 20% Mini-Protean TGX gels (Bio-Rad, number 456-1096). Gels were run at 80 to 100 V for 45 min to 1 h and transferred to Immobilon-FL transfer membranes (IPFL00010) using semidry transfer at 10 V for 45 min. Membranes were blocked in 5% bovine serum albumin fat-free milk prepared in PBS containing 0.05% Tween 20 for a minimum of 2 hrs at room temperature. Membranes were incubated with primary antibodies (anti-ARF6 antibody (3A-1), Santa Cruz, cat nr. sc-7971 and anti-β–actin, 8H10D10, Cell Signaling, cat nr. 3700S) at 4°C by rocking overnight. Antibodies were washed off the membranes with four 15-min rocking periods using fresh changes of PBS. Secondary antibody incubation was done using fluorescent LI-COR antibodies (anti-rabbit IgG, cat. nr. 926-68071; and anti-mouse IgG, cat nr. 926-32210) or regular secondary antibody (anti-mouse IgG HRP, Thermo Fisher, A9044) diluted 1:10,000 in PBS for 1 hr at room temperature with gentle rocking. Membranes were washed with PBS as before and visualized using a LI-COR Odyssey Imager (Huh7 cells) or a regular imager after development with the Pierce^™^ ECL Plus Western Blotting Substrate (Thermo Fisher, cat nr. 32132X3) (Calu-3 cells).

### Viability assay

Huh-7 cells or Calu-3 were seeded in a 96-well plate at 10,000 cells/well and allowed to adhere overnight. The next day, medium was replaced by medium with compounds or vehicle (DMSO, at the highest concentration used) and resazurin (Biotium, 30025-1) was added according to the manufacturer’s instruction. One day post treatment (to mimic infection condition), absorbance (590 nm) was measured with a Synergy^™^ HTX plate reader.

## Notes

### Competing Interest Statement

The authors have declared no competing interest.

### Summary of Updates

Title updated; Figure 4 has been updated to add the control for bound virus and remove the data on AA147; Figure S1 and S2 were modified to add i) toxicity control of the compounds used in Calu3 cells and ii) western blot of Calu3 cells ARF6 KO

